# Optimal control of Multiple Myeloma assuming drug evasion and off-target effects

**DOI:** 10.1101/2024.06.06.597698

**Authors:** James G. Lefevre, Brodie A.J. Lawson, Pamela M. Burrage, Diane M. Donovan, Kevin Burrage

## Abstract

Multiple Myeloma (MM) is a plasma cell cancer that occurs in the bone marrow. A leading treatment for MM is the monoclonal antibody Daratumumab, targeting the CD38 receptor, which is highly overexpressed in myeloma cells. In this work we model drug evasion via loss of CD38 expression, which is a proposed mechanism of resistance to Daratumumab treatment. We develop an ODE model that includes drug evasion via two mechanisms: a direct effect in which CD38 expression is lost without cell death in response to Daratumumab, and an indirect effect in which CD38 expression switches on and off in the cancer cells; myeloma cells that do not express CD38 have lower fitness but are shielded from the drug action. The model also incorporates competition with healthy cells, death of healthy cells due to off-target drug effects, and a Michaelis-Menten type immune response. Using optimal control theory, we study the effect of the drug evasion mechanisms and the off-target drug effect on the optimal treatment regime. We identify a general increase in treatment duration and costs, with varying patterns of response for the different controlling parameters. Several distinct optimal treatment regimes are identified within the parameter space.

**Author summary:** In this work we investigate a model of Multiple Myeloma, a cancer of the bone marrow, and its treatment with the drug Daratumumab. The model incorporates proposed mechanisms by which the cancer evades Daratumumab by reduced expression of the receptor CD38, which is the drug target and normally abundent in the cancer cells. The model includes an off-target effect, meaning that the drug treatment destroys some healthy cells alongside the targeted cancer cells. Both mechanisms can reasonably be expected to reduce the efficacy of the drug. We investigate the model using optimal control methods, which are used to find the drug dose over time which best balances the financial and health costs of treatment against cancer persistence, according to a specified cost function. We show that this drug resistence and off-target effect prolongs the optimal treatment and increase the burden of both the disease and drug. We analyse the distinct effects of the controlling parameters on each of these costs factors as well as the time course, and identify conditions under which extended treatment is required, with either intermittant treatment or a steady reduced dose. Extended treatment may be indefinite or for a fixed period.

## 1 Introduction

Myeloma is a plasma cell cancer that occurs in the bone marrow. Myeloma cells typically form masses of cancerous tissue, and the disease is known as multiple myeloma (MM) when more than one mass is present. Myeloma can crowd out healthy marrow tissue, leading to a range of potential deficiencies, and invade and weaken bone. It may also cause damage via production of abnormal antibodies. A number of treatment options are available, although a complete cure has proved elusive [1], [2].

In general, myeloma cells are marked by very high CD38 expression, motivating the use of the monoclonal antibody Daratumumab (Dara), which effectively targets myeloma via several mechanisms [3]. However, CD38 is also expressed in a wide range of cell types, resulting in important and complex off-target effects [4]. Daratumumab is a leading treatment for MM, commonly sold under the brand name Darzalez. We note that several other drugs have been developed to treat myeloma, such as Elotuzumab [5], [6] and Lenalidomide [7], which can be used together in combination with the adjunct drug Dexamethasone as a combination treatment for refactory disease [8]. However, in this work we consider treatment using Dara only.

Myeloma develops tolerance to Daratumumab. The dynamics are not fully understood, although various mechanisms have been proposed and combination therapies and recurrent treatment have had clinical success [9]. In this work we focus on one known tolerance mechanism, in which myeloma cells evade Daratumumab via loss of CD38 expression. This may occur passively due to differential response to Dara treatment, leading to a relative increase in myeloma cells with low CD38 expression. There may also be a direct loss of expression (without cell death) in response to drug exposure, as has been shown to occur in red blood cells [10].

Using optimal control theory, we investigate how these drug escape and off-target effects impact on effective treatment protocols for Dara that balance the cost of treatment with the burden of disease. We find in general that these effects support a more prolonged treatment regime and drive higher overall costs, and we further investigate the connection between the specific dynamics and the total cost and duration of treatment. Notably, we find that with a linear cost function, the optimal drug dosage over time can have distinct functional forms depending on parameter values. An initial period of maximal dosage may be followed by lower level treatment at constant or reducing dose, possibly after a pause. In certain cases where a more prolonged or indefinite treatment is required, we find that a regular intermittant treatment regime is optimal. This may help to inform maintainance Dara treatment, which has been shown to be effective in some cases [11].

### 1.1 Dynamical systems and optimal control theory

Dynamical systems are a class of mathematical model used to study complex time varying systems. The state of a system at any time is represented by one or more numerical *state variables*, and the rate of change of each state variable at a given time is taken to be a function of itself and the other state variables. These functions typically form a system of ordinary differential equations (ODEs), which can be solved or analysed using a range of numerical and analytical techniques, in order to provide insight into the modelled system. First applied by Poincaré to the study of the three body problem in classical mechanics [12], dynamical systems theory has been developed and applied extensively in a wide range of areas. Biological applications were pioneered with the logistic model of Verhulst [13], [14], representing exponential population increase constrained by a maximum carrying capacity. The famous Lotka–Volterra predator-prey model was first used to study interacting chemical species [15], then later applied to an ecological model, showing that the interactions between a prey species and a predator species could produce a continuing oscillation of populations over time [16].

Cancer biology typically involves complex interactions of cancer cells with their microenvironment and with a range of immune and other cell types, and a range of dynamical systems models have been developed to help understand this clinically critical biology [17]. State variables represent populations of cells and other relevant species. A number of papers (e.g. [18], [19]) have modelled cancer-immune interactions through a predator-prey framing, with cancer cells as the prey and cytotoxic T-cells, a type of white blood cell which destroy diseased cells, acting as the predator. Modelling of this interaction is of particular interest due to the introduction of CAR T-cell therapy [20], which relies on modified T-cells with an increased capacity to target cancer. A limitation of the predator-prey analogy is that consumption of prey strengthens the predator, whereas in cancer the first-order effect of immune cell “predation” weakens the immune cell population. However, positive feedback may be produced via various second order effects. The appropriate approaches for modelling these complex interactions is the subject of active research [21], [22].

Optimal control theory is used to study external interventions in a dynamical system. The *control* is an exogenous variable representing an external force. It is incorporated into the state equations, so that it may influence the rate of change of the state variables. The general form of the ODE system is then

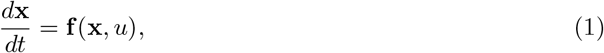

where **x**(*t*) is the vector of state variables and *u*(*t*) is the control. A cost function is defined based on the values of the control and the state variables over a specified time window [*t*_0_, *t*_*f*_], and the *optimal control* is chosen to minimise this cost function for a given initial state:

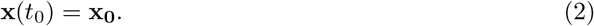

Since the control may vary freely over the time window, determining the optimal control is a challenging problem in general. Specialised numerical methods are required, with mathematical and numerical constraints that restrict the form of the cost function. Our approach is based on Pontryagin’s maximum principle [23]. The cost function must have the form

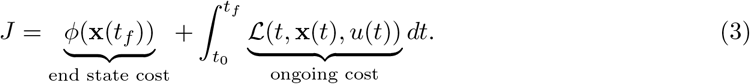

Cost functions in which ℒ has a linear dependancy on *u* require bounds to be imposed on *u* in order to give a well defined solution. The optimal control will generally take the form of a step function, equal to either the lower or upper bound at each time. The bounds typically correspond to “off” and “on”, and this is known as a *bang-bang* control. If ℒ is a convex function of *u* the problem is more tractable, generally giving a smoothly varying optimal control without the need to impose bounds. This is known as a continuous control. We will consider cost functions of both types (see Section 1.3).

Optimal control theory has been applied to clinical models to find theoretically optimal treatment regimes, with controls representing drug dose levels over time and cost functions designed to balance the monetary and health cost of treatment against the burden of disease. Important recent work includes applications to cancer immunotherapy, including generalised Lotka-Volterra predator-prey models [24] and models of combination therapy [25].

In this work we take a different approach, incorporating a simple immune response and placing focus instead on the drug escape mechanism and off-target effects discussed above.

### 1.2 Model of myeloma and Daratumumab

Crowell et. al. proposed a dynamical system model of blood cancer (ASL) incorporating a competition between healthy and cancerous cells for space in the marrow, with proliferation of both populations restricted as the total cell population approaches the carrying capacity [26]. The model features the migration of healthy cells into the compartment from a separate stem cell compartment, and migration of both healthy and cancerous cells into the blood system.

Sharp et al. [27] applied an optimal control methodology to a modified version of the Crowell model, with the addition of an immune response to cancer. The immune response was represented using a Michaelis-Menten term, which models a bounded immune capacity that initially scales with the cancer level but has a maximum capacity to remove cancer calls; this has the effect, for appropriate parameter choices, of allowing stable steady states with and without cancer present. This modification allows finite term treatment to result in a permanent control of the cancer; the authors found that this property was required for convergence of the optimal control algorithm. These works provide a calibrated model that supports the expected dynamics of cancer and cancer treatment, as well as a proven methodology for obtaining continuous and bang-bang controls.

We develop a dynamical systems model of myeloma that adapts the core features of the Sharp model. Our model also contains healthy and cancerous populations of marrow cells that compete for space, while the downsteam blood cell populations are dropped, as they do not affect the marrow cells or the cost function. The upsteam stem cell population is modelled implicitly as an influx of healthy cells into the marrow (rate *β*_*A*_) under the assumption that the stem cell population is at steady state; this causes only a transitory divergence from the Sharp model.

In order to incorporate the core drug escape mechanism, we replace the cancer population with CD38+ and CD38-cancer cell subpopulations *P* and *N*, of which only *P* is susceptible to the drug. These represent alternate states of a single population, so it is assumed that cells move between *P* and *N* at rates *δ*_*P*_ and *δ*_*N*_. We will refer to this mechanism as expression switching. Given the general overexpression of *CD*38 in myeloma, we assume that *δ*_*N*_ = 10δ_*P*_, and that fitness is significantly lower in *N*. We also allow for both direct drug-induced loss of CD38 expression (*δ*_*P u*_), and an off-target effect modelled by drug induced death of healthy cells (*µ*_*Au*_). Note that off-target drug effects can be modelled implicitly in the cost function, but this explicit approach accounts for interaction with population dynamics.

The complete model is as follows, where the three state variables **x** = (*A, P, N*) are each expresed as a proportion of the marrow carrying capacity:

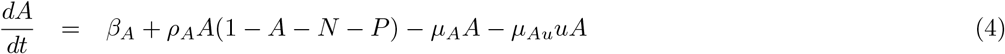

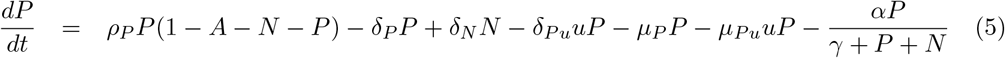

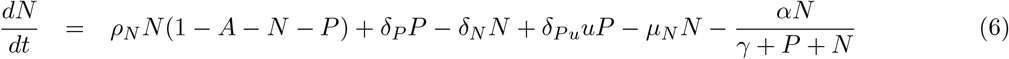

Here the control *u* ≥ 0 represents the dosage rate of Daratumumab. The state variables must also be non-negative to be physically realisable. The model parameters, and the default values used, are listed below.

| Description | Parameter | Value |
| --- | --- | --- |
| Influx of healthy cells | $\beta_A$ | 0.1008 |
| Proliferation rate of $A$ (healthy cells) | $\rho_A$ | 0.43 |
| Rate of death or other exit for $A$ | $\mu_A$ | 0.44 |
| Off-target mortality effect on $A$ per unit of Dara (control) | $\mu_{Au}$ | 0.1 |
| Proliferation rate of $P$ (CD38+ myeloma) | $\rho_P$ | 0.28 |
| Rate of death or other exit for $P$ | $\mu_P$ | 0.048 |
| Additional death rate of $P$ per unit of Dara (control) | $\mu_{Pu}$ | 1 |
| Proliferation rate of $N$ (CD38- myeloma) | $\rho_N$ | 0.15 |
| Rate of death or other exit for $N$ | $\mu_N$ | 0.06 |
| Rate of loss in CD38 expression in $P$ | $\delta_P$ | 0.003 |
| Rate of gain in CD38 expression in $N$ | $\delta_N$ | 0.03 |
| Increased loss in CD38 expression in $P$ per unit of Dara | $\delta_{Pu}$ | 0.2 |
| Immune control rate | $\alpha$ | 0.015 |
| Immune control half saturation | $\gamma$ | 0.1 |

Where possible, parameter values were adapted from the Sharp model, which were selected to produce balanced dynamics supporting both healthy and cancerous states and the capacity for effective drug control. Proliferation and exit rates are set so that the CD38+ myeloma cell population *P* is slightly more fit than the cancer population in the Sharp model, and the CD38-population *N* substantially less fit. The effect of a unit of control on the mortality rate of CD38+ cancer cells, *µ*_*P u*_, is fixed at one; this defines the scale for the control *u*. Since CD38 is typically highly overexpressed in myeloma, it can be assumed that *µ*_*Au*_ is substantially lower than *µ*_*P u*_ = 1; we use 0.1 by default, although higher values are also considered. We also choose a conservative initial value of *δ*_*P u*_ = 0.2, implying the direct loss of expression from Dara is a smaller effect than mortality, but with higher values considered. Note that in our model, as in the Sharp and Crowell models, the unit of time is abstract and parameters are not calibrated to real data.

### 1.3 Cost functions and control types

An optimal control can only be calculated in reference to a cost function. This function encodes the health cost of cancer presence, as well as the cost of the drug dose over time — both its direct financial cost and health effects due to its side effects. However, the most appropriate mapping between these factors and cost is not obvious, including the correct weighting between cancer and drug dose.

Since results will depend on the cost function chosen, we consider two optimal control cost functions, corresponding to a continuous and a bang-bang control, to provide insight into the influence of the cost assumptions and the robustness of any conclusions. In each case the cost function takes the form of (3) with *ϕ*(**x**(*t*_*f*_)) = 0; removing the dependence on the final state is generally preferred as it provides more tractable computations. The health cost due to cancer is assumed to depend on the total cancer population, *P* + *N*.

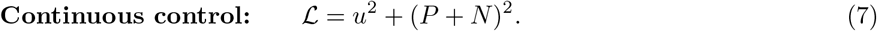

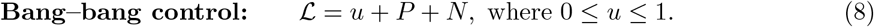

While it is normally expected that this second cost function will result in convergence to a control solution which is equal to either *u*(*t*) = 0 or *u*(*t*) = 1 for each value of *t*, the iterative update algorithm used means that solutions of this form are not guaranteed. Valid alternative forms were found in some cases, and a modifed form of the iterative update algorithm was used to improve convergence in these cases; see Methods 4.3 for details.

Note that the cost function cannot take into account the system state outside the selected time window, such as a cancer recurrence, and this must be taken into account in the interpretation of results. For example, the algorithm can return a null optimal control, with *u*(*t*) = 0 for all *t*, but this may be due to the time window being too short.

## 2 Results

### 2.1 Drug escape mechanism produces expected resistance to control

Our model extends the Sharp model [27], which we use as a negative control to validate the drug escape and off-target mortality effects. We can reproduce the core features of the Sharp model by suppressing the drug resistant CD38-cancer cell population (N) and the off-target drug effect. We verified this by defining a Null-N model with the parameter changes *N* (0) = 0, *δ*_*P*_ = 0, *δ*_*P u*_ = 0, *µ*_*u*_ = 0, *ρ*_*P*_ = 0.27, *µ*_*P*_ = 0.05. Simulations using this model are shown in Fig. 2 a-d. Without treatment, the healthy and cancerous cells reach a balance. The presense of the immune response shifts this balance against the cancer without achieving elimination. But if the treatment can reduce the cancer level sufficiently, the immune response will prevent recurrence.

**Figure 1.**
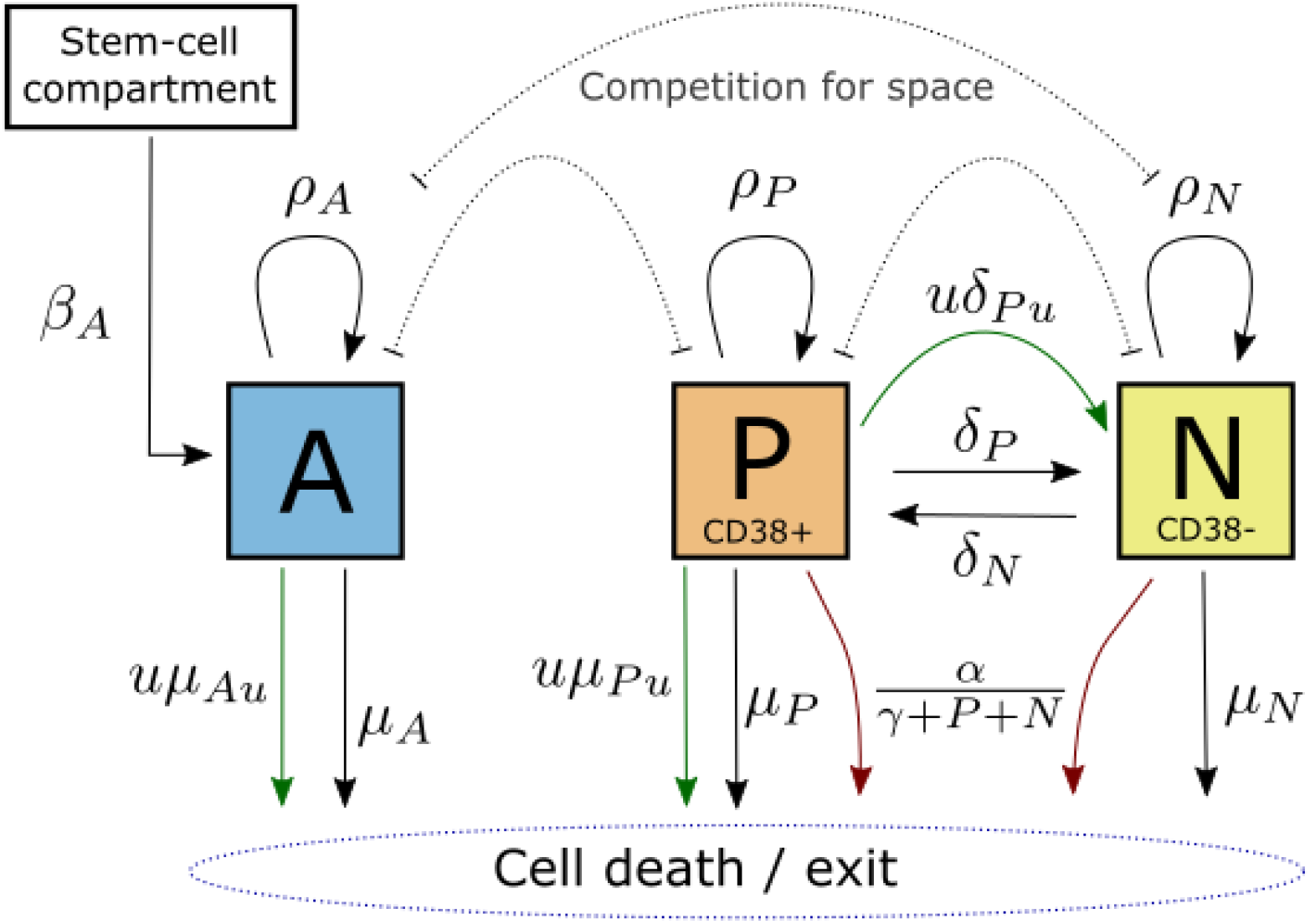
Model of multiple myeloma (MM) treatment with Daratumumab (control, *u*), including an immune response (red arrows) and drug escape mechanisms via loss of CD38 expression. Three drug actions are included (green arrows): cell mortality and loss of CD38 expression in the CD38+ cancer cells, and off-target cell mortality in healthy cells within the compartment.

**Figure 2.**
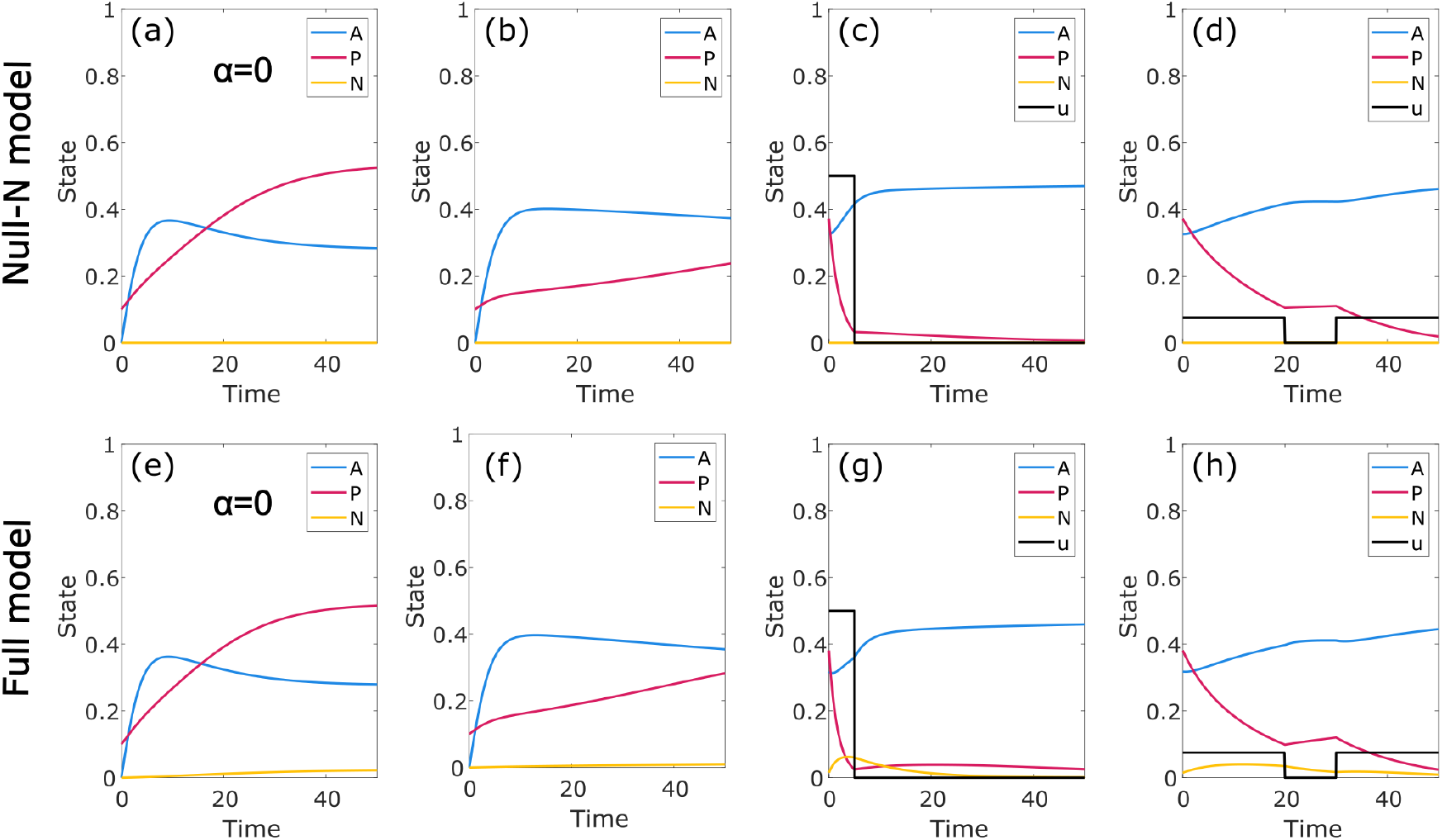
Model validation and comparison with Sharp model. Selected numerical simulations using the fourth order Runge-Kutta method and time step 0.001. The full model developed in this paper (e-h) is compared with a simplified version designed to replicate the Sharp model (Null-N model, a-d), in which the CD38-cancer population is suppressed. In a,b,e,f the initial state is *P* = 0.1, *A* = *N* = 0 and no control is applied; in a,e we also suppress the immune response, as in Sharp Figure 2. In c,d,g,h the simulation starts at steady state and a prespecified control is applied.

In Fig. 2 e-h we show the corresponding simulations using our full model. As intended, in the absense of the drug control the CD38-population plays only a marginal role; this can be seen in e,f. In the presense of the drug control, the role of this population grows and has the effect of diminishing drug efficacy via the escape mechanism. Panels g,h suggest that the drug escape mechanism plays a larger role when the control has higher intensity and shorter duration; we examine this issue more systematically below through the lens of optimal control. For the parameters used here, where the drug’s effect on healthy cells is only one tenth of its effect on CD38+ cells, the off-target drug effect is extremely minor.

### 2.2 Drug escape motivates prolonged treatment

Using the continuous and bang-bang optimal controls, we can evaluate the overall effect of the model modifications under the default parameters in terms of the cost to treat and optimal pattern of treatment (Fig. 3). Note that cost is not comparable between the two types of optimal control. In both cases, the total control and overall cost is increased relative to the Null-N control, and the duration of treatment is extended. For both control types a high initial drug dose in the full model rapidly reduces overall cancer levels, but at the cost of much higher levels of the drug-immune CD38-population, and recovery of the healthy cell population is inhibited by the off-target effect while the control dose is high. The control is then continued at a lower level that balances these factors, until control of the cancer is achieved.

**Figure 3.**
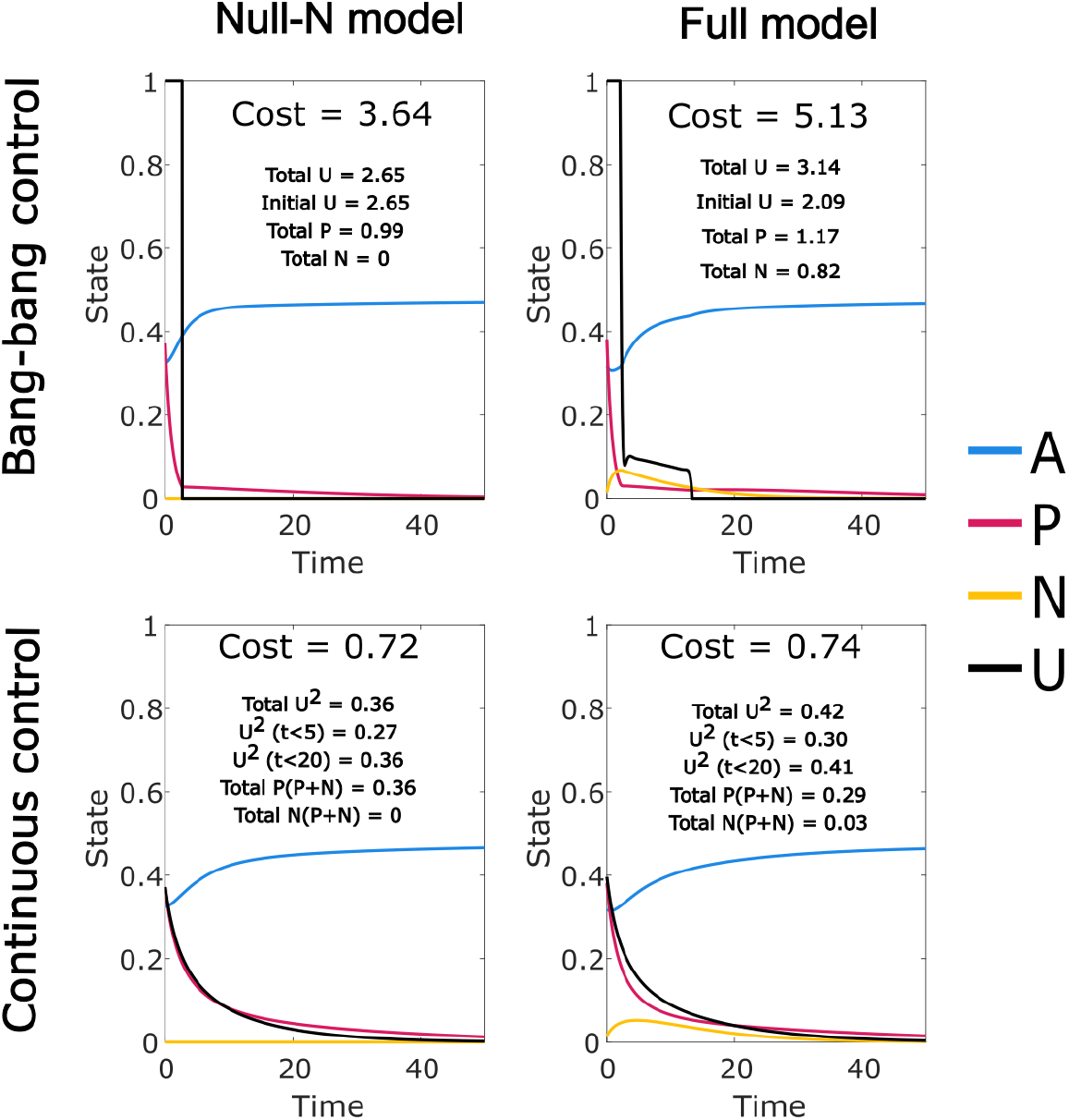
Optimal controls for the full model developed in this paper (Full model) and the simplified version that replicates the Sharp model (Null-N). In each case we give the overall cost function value ((7) or (8)) and its components (control cost and cancer burden). We also note the total control applied in the initial period when the control is at its maximum level. For the continuous control cost function, the total cancer cost ((*P* + *N*)^2^) is allocated proportionately between *P* and *N* for the quoted numbers. The drug related cost incurred in the first 5 and 20 time units is also given as an indication of relative control duration. Optimal controls were found using a time period of length 200, plots and numerical results are shown for the initial 50.

The increases in cost, total control, and duration of treatment relative to the Null-N control are all much smaller for the continuous control than for the bang-bang control. This can likely be attributed to the fact that prolonged, low level treatment is favoured by the quadratic function (7), incurring a low cost. This explains the tapered shape of the continuous control solutions for the Null-N model, and with the addition of the new mechanisms the required prolongation of treatment is small and achieved at low cost. However, the quadratic cost will not reflect the financial cost of drug supply or treatment, and the very low cost associated with prolonged low level treatment may not be realistic.

In contrast, the linear cost function used in the bang-bang control does not discount the cost of continuing lower level treatment. This cost function typically gives optimal control levels at either the maximum level or zero; in the Sharp model all solutions consisted of an initial period at maximum level followed by an abrupt and final end of treatment. The fact that in our model the bang-bang optimal control includes a period of intermediate level control provides clear support for extended treatment.

### 2.3 Reduced immune response requires extended treatment regime

The immune response, controlled by parameter *α*, plays an important role in treatment. In Fig. 4 we show the affect of varying this parameter on the optimal control. We see that a reduced immune response increases the cost to treat primarily through prolongation of the control; for the bang-bang control, the initial period of treatment is almost invariant. As the immune response approaches zero, there is a point at which final control of the cancer becomes impossible, and the optimal control transitions to an initial high dose treatment followed by an indefinite maintainance treatment. At this stage the burden of CD38-cancer becomes significant; we can also project ongoing costs from the trendline.

**Figure 4.**
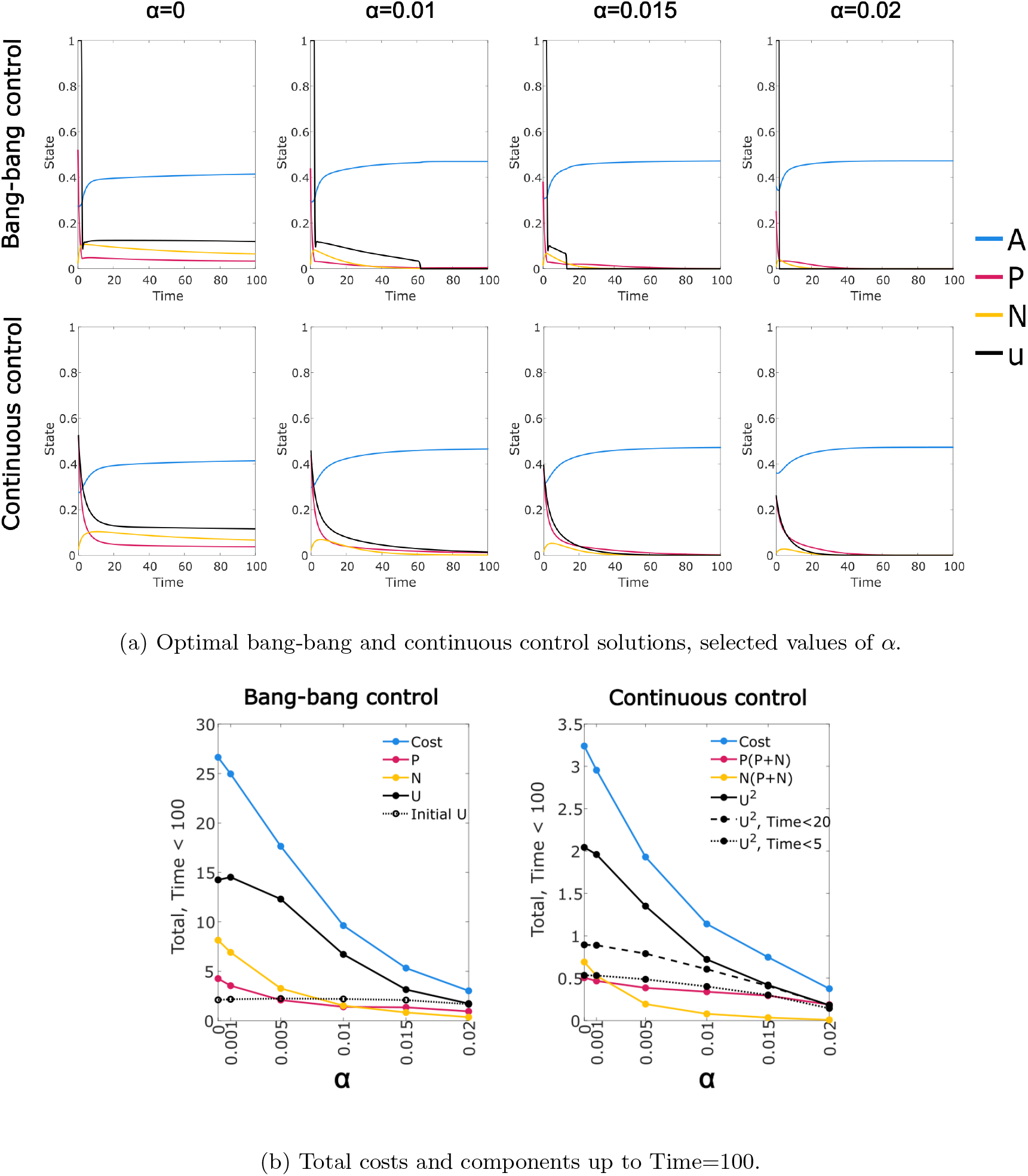
Optimal control solutions for range of *α* values (original value *α* = 0.015).

We are most interested in model parameters that allow both a persistent cancer state and the possibility of permanent control via drug treatment. We show in Section 4.4 that this requires the Michaelis-Menten immune response; a linear immune response can be regarded as a simple modification of the exit rate parameters and cannot achieve the same effect. However, cases in which the cancer must be managed through ongoing treatment are also of interest, despite posing some difficulty in interpretation due to the finite time window used in the optimal control methodology. From a modelling perspective, it is significant that when the immune response is not sufficient to allow permanent control of the cancer, our algorithm is able to find an optimal steady state treatment regime, as shown in Fig. 4 when *α* = 0. This can be attributed to the additional mechanisms in our model, as it is not the case for the Null-N model. If we consider the Null-N model with a constant level of control applied so that *P* approaches 0, then *A* will approach a steady state *A*_0_ and we have 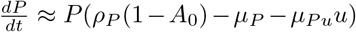, giving an asyptotically exponential solution for *P*. Cessation of the control will lead to exponential increase until *P* is again non-negligible. This implicitly models cancer at arbitrarily low levels, potentially less than a single cell, and also results in convergence failure of the optimal control, due to extreme insensitivity to control timing while *P* is at a negligible level. Thus we see that our modified model provides an improvement in this case both computationally and as a biologically realistic system.

### 2.4 Drug escape parameters have distinct and interacting effects on model dynamics

The drug escape mechanism we model consists of four added features: an alternative CD38-cancer state with reduced fitness; switching of cancer cells between the CD38+ and CD38-states; a response to treatment in the form of loss of CD38 expression; and mortality of CD38+ but not CD38-cancer cells in response to the control. Here we consider the sensitivity of the model and the optimal control solutions to the parameters controlling these features. The mortality effect is kept fixed with *µ*_*P u*_ = 1 while we consider variations in the other three features, observing distinct responses in each case.

For the expression switching mechanism, in which cancer cells lose or gain CD38 expression, we retain *δ*_*N*_ */δ*_*P*_ = 10 to reflect the typical dominance of the CD38+ state, but consider large coordinated changes in both values (Fig. 5). There is minimal effect on the optimal continuous control. For the bang-bang control, higher rates of switching lead to a prolonged optimal control, with a control that is higher in aggregate despite a shorter initial period at maximum intensity. The temporal pattern of the control after the initial period also changes: at the highest level of expression switching there is a much stronger reduction over time in the control level. The aggregate cancer level remains relatively constant, although CD38 expression increases.

**Figure 5.**
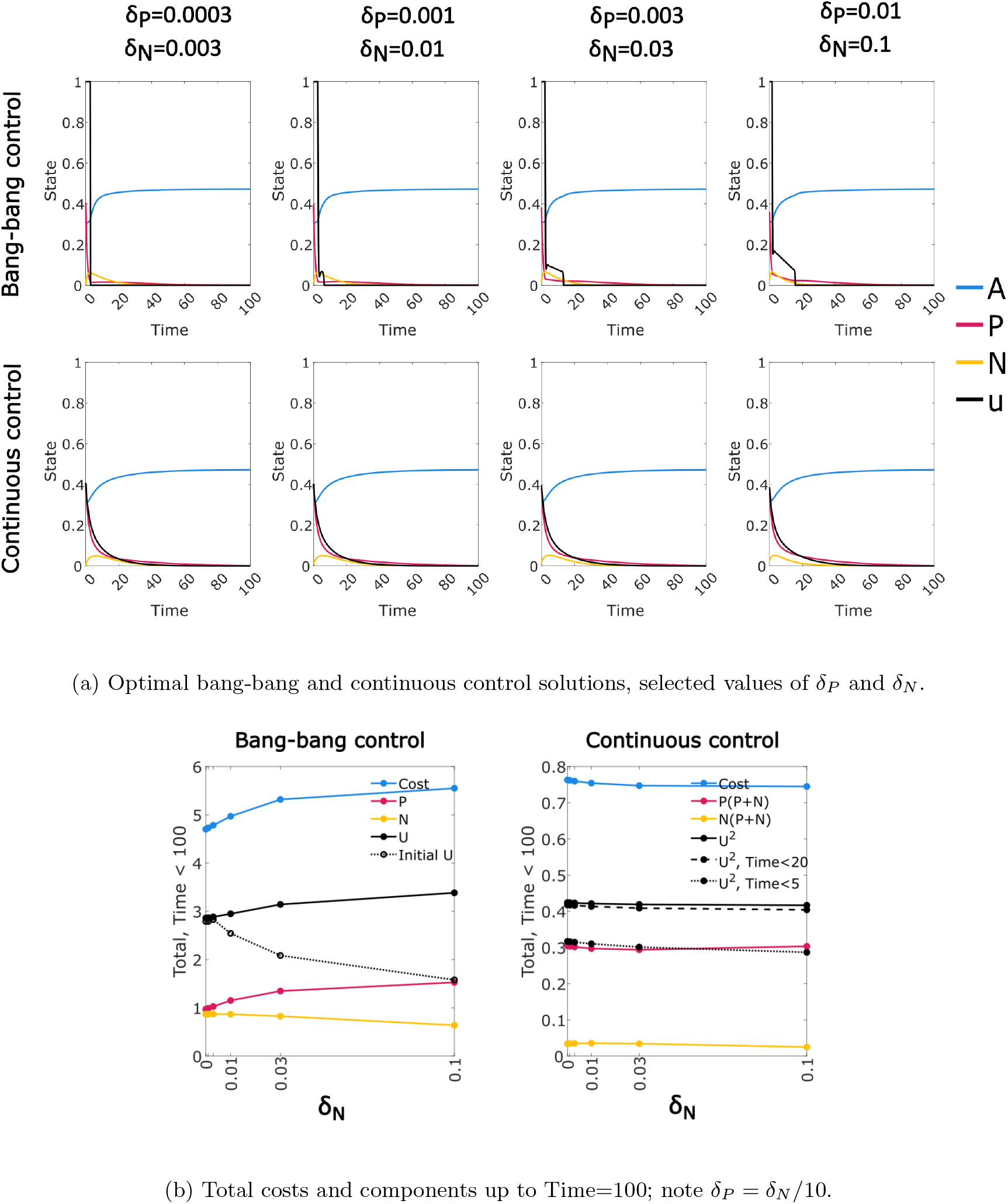
Effect of expression switching of CD38 expression on optimal treatment: optimal control solutions for range of *δ*_*P*_ and *δ*_*N*_ values (original values *δ*_*P*_ = 0.003, *δ*_*N*_ = 0.03).

The fitness penalty from loss of CD38 expression in cancer cells is represented by a reduction in proliferation and increase in mortality. Plausible variations in the size of this penalty have little effect on outcomes except for a modest reduction in treatment duration at higher fitness penalty (Fig. 6). However, if the fitness penalty is removed entirely (left side) there is a large increase in cost driven by extended control and persistent CD38-cancer cells. Note that we do not attempt to disentagle the effects of changes in proliferation and mortality.

**Figure 6.**
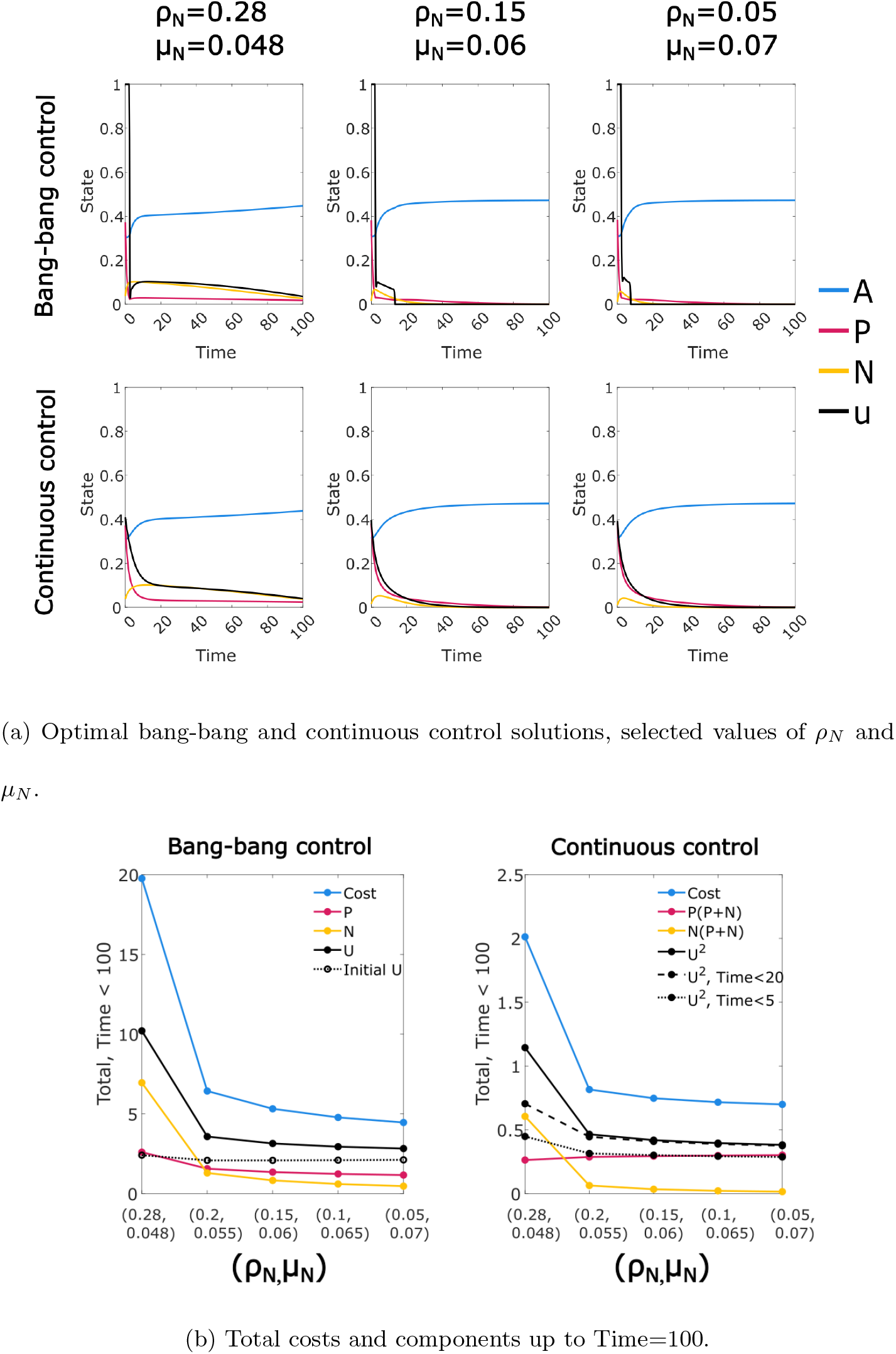
Effect of CD38-myeloma fitness on optimal treatment: optimal control solutions for range of *ρ*_*N*_ and *µ*_*N*_ values. The *x* axis represents fitness of the CD38-cancer cells, with fitness decreasing to the right, and the leftmost value representing fitness equal to the CD38+ cells (original value *ρ*_*N*_ = 0.15 and *µ*_*N*_ = 0.06).

Finally, the most complex response is elicited from variation in the rate of drug-induced loss of CD38 expression (Fig. 7). Optimal control solutions appear to show competing effects from this loss of CD38 expression: the drug control induces a CD38-subpopulation that persists under treatment, imposing a health burden and requiring more prolonged treatment (particularly for the bang-bang control). However, the lower fitness of this subpopulation results in a relatively stable or reduced quantity of control required in aggregate.

**Figure 7.**
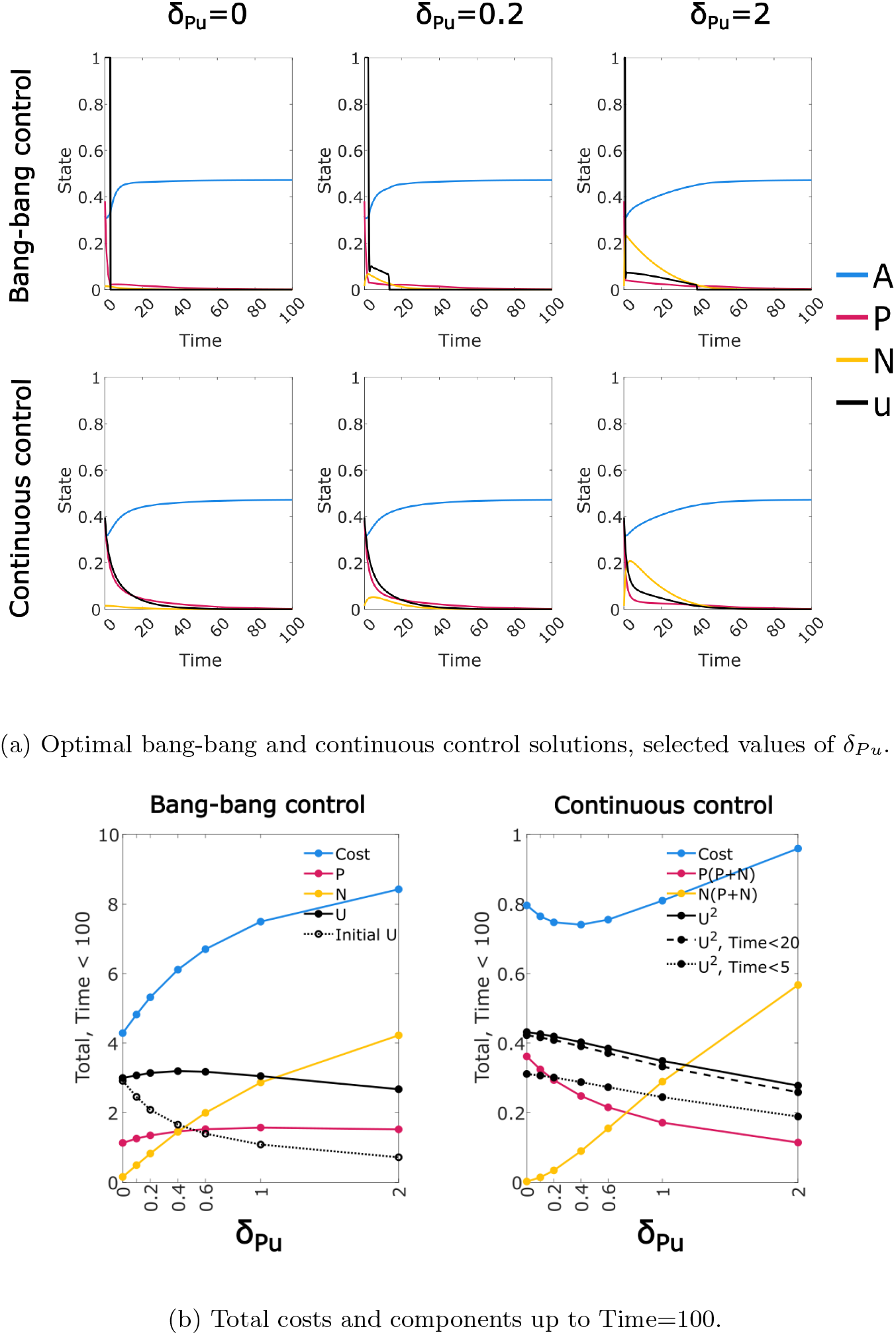
Effect of drug-induced loss of CD38 expression on optimal treatment: optimal control solutions for range of *δ*_*P u*_ values (original value *δ*_*P u*_ = 0.2).

The most striking feature we observe from the optimal control analysis of our model is the prolongation of treatment at lower intensity in the bang-bang control, which is significant precisely because it is not typically present in bang-bang controls. Note that while a modified method was required to obtain these solutions, these solutions are not an artefact of the method, as detailed in Methods 4.3. We see that the existence of this phenomenon requires both the induced loss of CD38 expression and expression switching between CD38+ and CD38-states.

### 2.5 Elevated off-target drug effect produces distinct form of bang-bang control

In addition to modelling drug avoidance via loss of CD38 expression, our model includes an off-target effect, in which the control causes some mortality in healthy cells. While the harm caused by drug side-effects can be modelled through the cost function, including this feature explicitly allows us to examine the effects on the population dynamics, and particularly the interaction with the drug escape mechanism. Since myeloma cells notably over-express the drug target CD38, we expect realistic values of the off-target mortality parameter *µ*_*Au*_ to be much less than 1. At this level the off-target effect appears to have limited influence. When we consider higher values (Fig. 8) we see a general pattern of slightly increased costs (both drug and disease burden). However, at *µ*_*Au*_ = 1 we see a striking change in the form of the optimal bang-bang control. Instead of a period of continuing control at reduced intensity, the initial period of maximum intensity control is followed by a complete cessation of treatment, then a second shorter period of maximum intensity treatment. During the break in treatment the healthy cell population recovers while the drug resistant CD38-population declines, but the CD38+ cancer subpopulation also recovers from low levels. The followup treatment prevents a cancer resurgence, reducing levels to where they can be controlled by the immune response.

**Figure 8.**
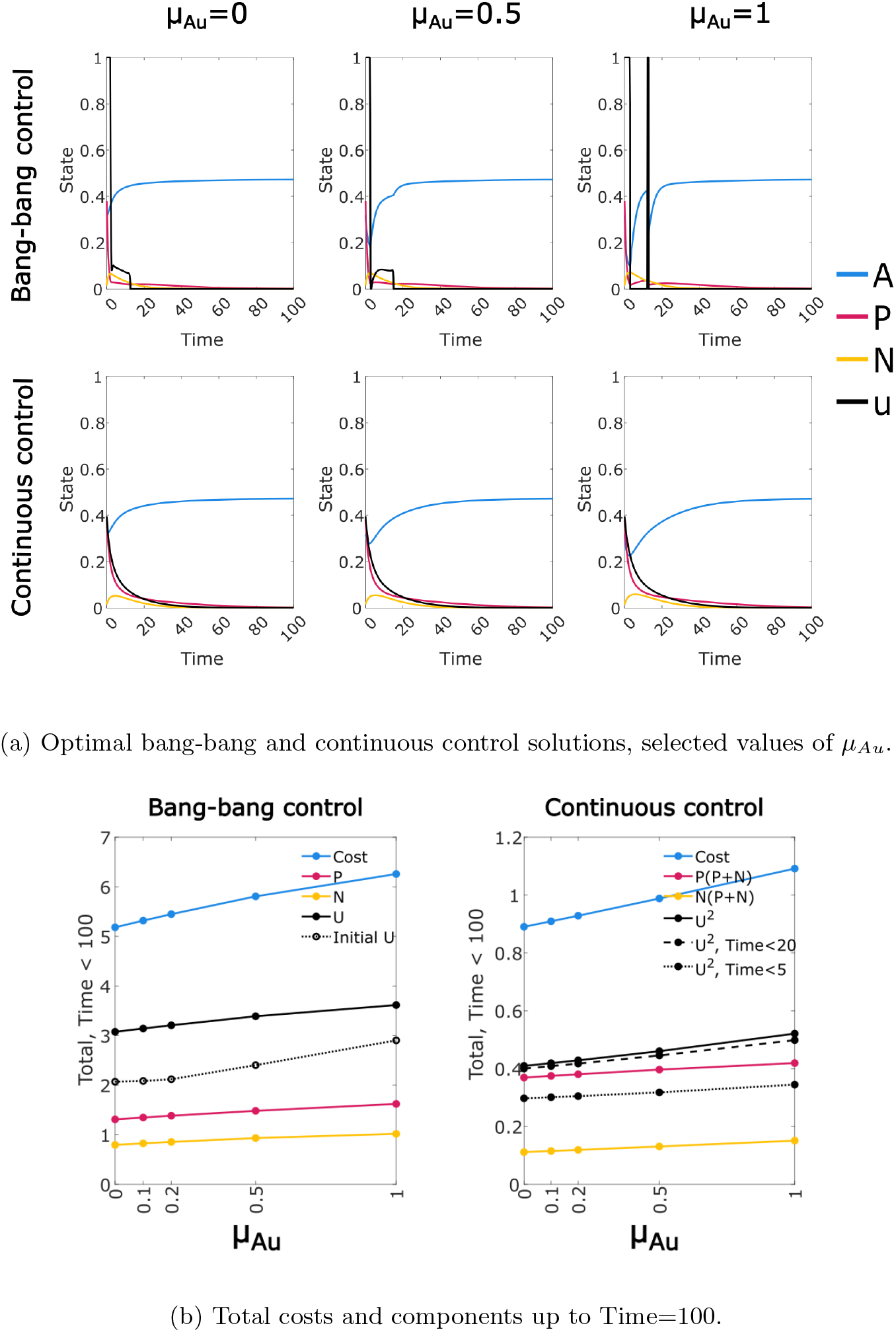
Optimal control solutions for range of *µ*_*Au*_ values (original value *µ*_*Au*_ = 0.1).

### 2.6 Optimal bang bang controls may be cyclic or discontinuous

The bang-bang optimal control solution with *µ*_*Au*_ = 1 raises the question of whether the solution may take other forms depending on the choice of parameters, particularly cyclic or discontinuous control solutions. The value *µ*_*Au*_ = 1 seems biologically implausible, so we performed a systematic search for alternate forms of the optimal bang-bang control using a somewhat more reasonable value *µ*_*Au*_ = 0.5. The rate of expression switching appeared to influence the shape of the control, so we considered both 10-fold increase and 10-fold decrease in these values (maintaining a fixed ratio between them). Drug-induced loss of CD38 expression also plays a key role, so we considered the effect of a 10-fold increase in this rate (*δ*_*P u*_). In addition, we considered removal or reduction in the immune intensity (*α* = 0, 0.01 instead of *α* = 0.015).

We see a remarkable solution form in the case of reduced expression switching, with reduced or zero immune response (Fig. 9, top row). The optimal control takes the form of short periods of control at maximum intensity, separated by longer periods of zero control. The initial treatment period is longer, and is also followed by a longer break; the treatment periods then follow a regular pattern. When the immune response is removed (*α* = 0), permanent control of the cancer is not possible, and the solution tends towards a repeating cyclic pattern. When *α* = 0.01 we see a modifed version of this pattern which terminates when suppression by the immune response has been established. Increasing the rate of drug-induced loss of CD38 expression appears to suppress this cyclic solution (Fig. 9, bottom row), however we retain a period of zero control following the initial period of maximum-intensity control. Note that the cases with unchanged or increased rates of expression switching (*δ*_*P*_ and *δ*_*N*_) did not give any cyclic or discontinuous solutions (data not shown).

**Figure 9.**
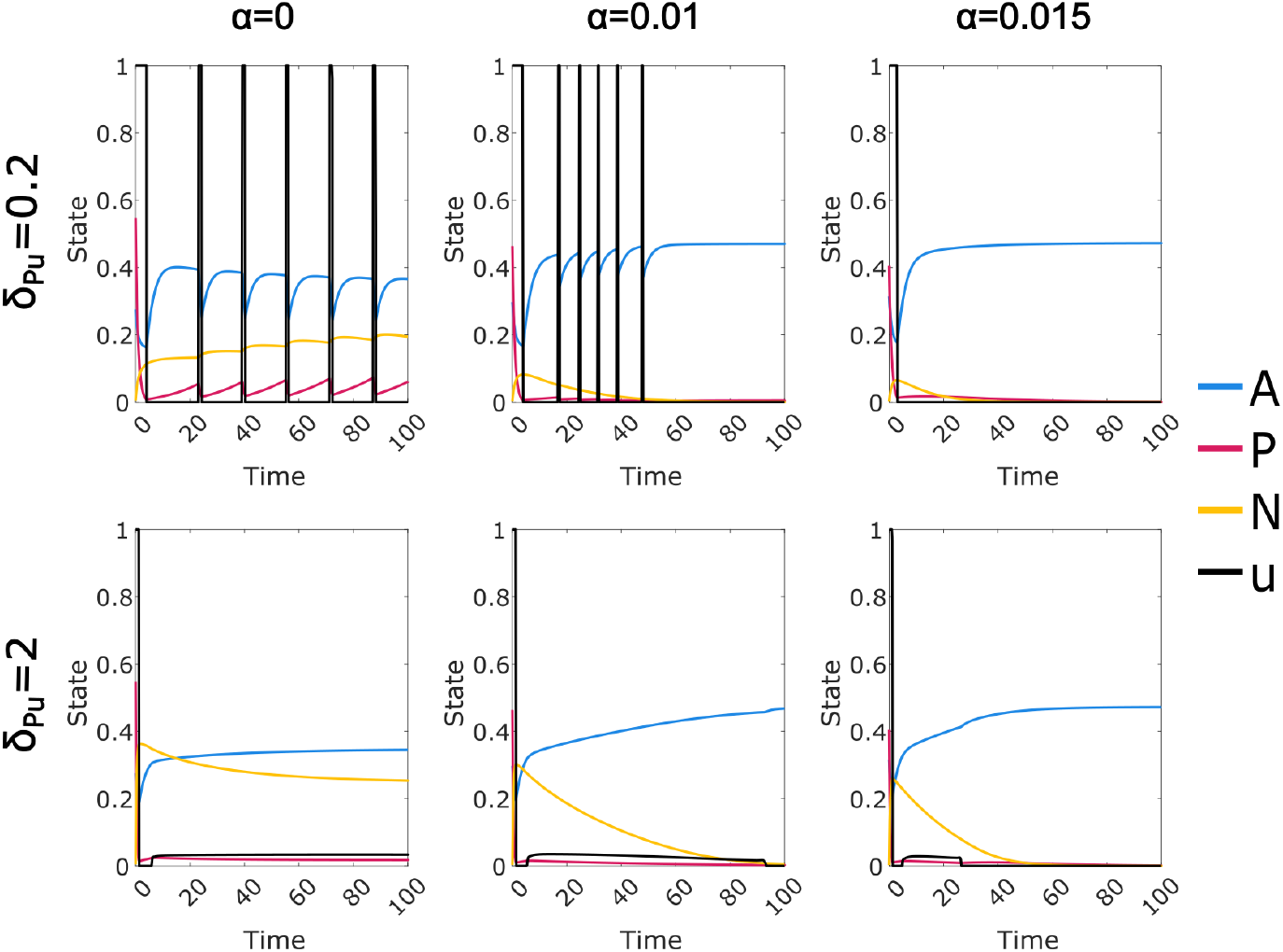
Optimal bang-bang control solutions for *µ*_*Au*_ = 0.5, *δ*_*P*_ = 0.0003, *δ*_*N*_ = 0.003, *α* = 0, 0.01, 0.015, *δ*_*P u*_ = 0.2, 2.

Since we see the cyclic solutions (Fig. 9) at the lowest values of *δ*_*P*_, *δ*_*N*_ and *δ*_*P u*_ that were considered in this experiment, it is natural to ask whether the complete removal of one or both of these features would also give optimal control solutions with a cyclic form. We retain *µ*_*Au*_ = 0.5 and set *α* = 0, consistent with the clearest examples of cyclic solutions seen. We observe (Fig. 10) that the cyclic solution form appears to be consistent with *δ*_*P u*_ = 0, but not with *δ*_*P*_ = *δ*_*N*_ = 0. We have used a high value of the off-target mortality parameter *µ*_*Au*_ under the assumption that this is required to produce the cyclic solution form. We check this assumption by setting *µ*_*Au*_ = 0 in two cases with the clearest observed cyclic behaviour (Fig. 11), leading to a loss of the cyclic form.

**Figure 10.**
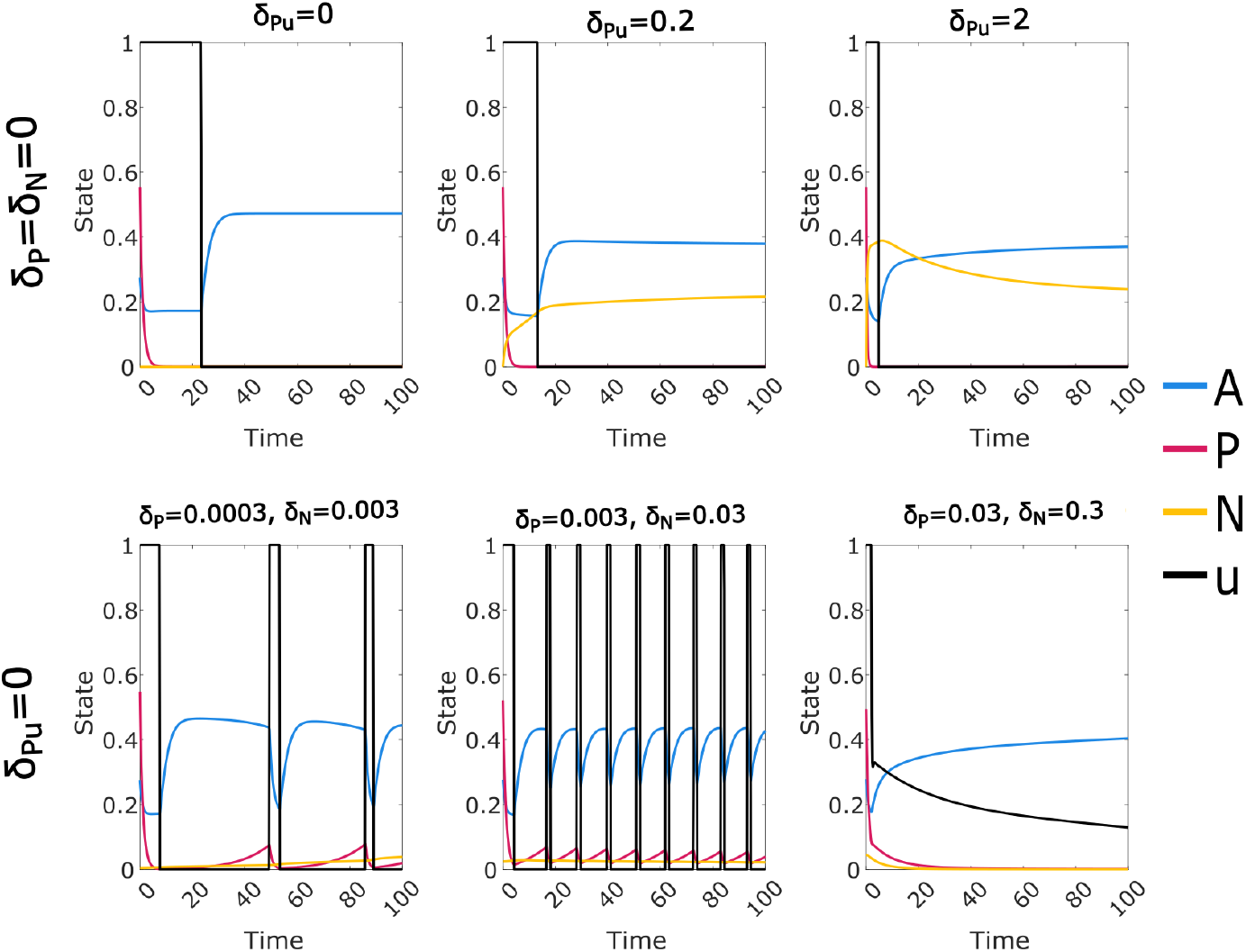
Optimal bang-bang control solutions with either *δ*_*P*_ = *δ*_*N*_ = 0 or *δ*_*P u*_ = 0; in all cases *µ*_*Au*_ = 0.5 and *α* = 0.

**Figure 11.**
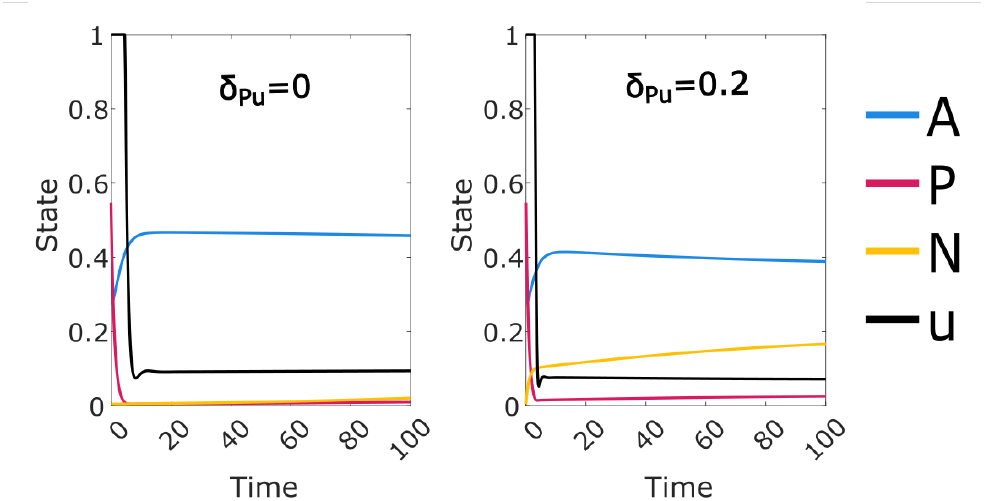
Optimal bang-bang control solutions for *µ*_*Au*_ = 0, *α* = 0, *δ*_*P*_ = 0.0003, *δ*_*N*_ = 0.003, and *δ*_*P u*_ = 0, 0.2.

While this analysis does not amount to a full exploration of the parameter space, our investigation suggests that the cyclic form requires a high value of *µ*_*Au*_, a low value of *α*, and a low but non-zero value of *δ*_*N*_ and *δ*_*P*_.

## 3 Discussion

We have presented a model of myeloma treatment using the monoclonal antibody Daratumumab, with which we investigated the impact of a drug escape mechanism and off-target cell mortality using optimal control theory. In our model myeloma cells evade the effect of Daratumumab via loss of CD38 expression, albeit at the cost of reduced fitness. This loss of expression may result from either differential mortality or as a direct result of drug exposure. The proposed mechanisms generally resulted in increased overall costs and extended duration of treatment. These mechanisms are modelled with several rate parameters, and in most cases the relationship between the rate parameters and outcomes such as total drug dose, treatment duration and cancer persistence were found to be at least directionally consistent between the two cost functions considered, suggesting that the identified trends are robust. Exceptions included the rate of expression switching (Fig. 6), which had very little effect on the continuous control, and the rate of drug-induced loss of CD38 expression, which had a somewhat inconsistent effect (Fig. 7). The forms of the optimal control solution over time presented a more complex situation, with greater differences between the two cost functions over a range of parameters.

The Null-N model, which we use as a negative control that reproduces the core Sharp model, gives optimal control solutions of two forms, depending on the cost function. The linear cost function with bounded control values gives solutions of the expected “bang-bang” form, in which the control starts at the maximum level and then at some time point permanently switches to zero. Intuitively, there is no advantage to delay in treatment, and so the total drug dose necessary to contain the cancer is administered in the minimal possible time. The quadratic cost function gives continuous solutions which begin at a high level then drop continuously, a tradeoff between removing cancer as quickly as possible and the cost advantage of treatment at lower dose.

When we included the drug escape and off-target effect mechanisms in the model, we found that for many parameter choices the bang-bang solution features a period of lower-intensity or intermittent treatment subsequent to the initial period of maximum level control. We can understand this as a period in which the imperative to treat the cancer must be balanced against the need to allow time for the CD38 expression level in the myeloma cells to recover, so that the drug effectiveness is restored. Recovery of the healthy cell population may also be a factor in this pattern.

The use of the quadratic cost function is motivated by the observation that the health burden of both disease and drug treatment will potentially increase at a super-linear rate; double the amount of drug or cancer causes more than twice the harm. However, this cost function also promotes extended treatment at very low dose, and the resulting tapering off of treatment appears to partly obscure the effect of the drug escape mechanism; the effect of our model modification is lower when using the quadratic cost function, particularly in terms of the prolongation of treatment. This tapering effect should be interpreted with appropriate caution in real world applications: below some level, the quadratic cost function will not fairly reflect the fixed financial cost of Daratumumab, or the practicalities of drug administration by diffusion. In contrast, for “bang-bang” control solutions using the linear cost function any period of ongoing control at a reduced level can be reasonably attributed to the biological mechanisms that we model.

Bang-bang controls for the full model take a range of forms, depending on the parameter values. These include the simple form with an initial period of maximum control and then no subsequent treatment. Higher levels of CD38 expression switching (*δ*_*P*_ and *δ*_*N*_) and drug-induced loss of CD38 expression (*δ*_*P u*_) produce controls with an intermediate period of ongoing control at a reduced level. Lower drug-induced loss of expression and lower but non-zero levels of expression switching, together with an elevated off-target effect (*µ*_*Au*_), tend to produce periodic control solutions with intermittant control at the maximum level. Both of these more complex forms are promoted by a reduced immune response. Insufficient immune response results in optimal control solutions in which indefinite continuation of treatment is required, either intermittant, or continuous at a reduced level.

We briefly note some limitations of this study. Although the identification of distinct optimal control forms depending on the various rate parameters is of considerable interest, this is principally theoretical. It demonstrates that real world optimal treatment regimes may be contingent on such factors, but the mechanisms discussed in [9], for example, are unquantified; the natural variation of CD38 expression between myeloma cells, and the rate at which this changes naturally and in the presense of Daratumumab, is unknown. We also did not attempt to definitively characterise the optimal control solutions across the entire plausible parameter space. This is largely due to the complexity of the system, but the reduction in convergence speed for the more complex bang-bang controls was also a limitation, and improvements to the convergence algorithm could allow a more complete analysis. Finally, a fully realistic cost function for a diffusion treatment such as Daratumumab would most likely incorporate a fixed per-session cost, reflecting factors such as setup and travel. This cannot be directly represented in the form of (3), and finding a method of incorporating such a cost factor could be of value.

### 3.1 Conclusion

In this work we investigated a drug evasion mechanism proposed by Saltarella et al [9], formalising the mechanism and incorporating it into a dynamical systems model of MM under Daratumumab treatment. Using simulations and optimal control methodology we validated the model, demonstrating that the proposed evasion mechanism can lead to effective resistance. We found a stronger resistance to higher drug dosage, resulting in an increase in both the cost and duration of treatment. We also demonstrated that our model is effective at representing disease under conditions in which a complete cure is impossible, and found optimal control solutions in these cases that include optimised ongoing treatment. Acknowledging the caveats discussed above, we also found several distinct functional forms for the optimal control across the plausible parameter space (using a linear cost function). This indicates that the optimal pattern of treatment may vary considerably depending on the cancer cell dynamics as well as patient characteristics such as the strength of immune response. While this analysis is theoretical, we have shown that the approach provides a promising framework for understanding this drug evasion mechanism in the case of Daratumumab in MM or any analogous system, and has the potential to inform empirical investigations leading to clinical advances.

## 4 Methods

### 4.1 Application of Pontryagin’s maximum principle

We begin by outlining the application of Pontryagin’s maximum (minimum) principle to solve an optimal control problem as specified in Section 1.1. Methods broadly follow [27] except for modifications to the convergence algorithm for the bang-bang control and to the method of obtaining equilibrium solutions.

Consider a boundary value problem of the form given by Equations (1) and (2), where **x**(*t*) = (*x*_1_(*t*), *x*_2_(*t*), …, *x*_*n*_(*t*)) is a vector of state variables and *u*(*t*) is the control. The objective is to choose *u*(*t*) to minimise a cost function of the form (3) over a time window [*t*_0_, *t*_*f*_]. We introduce a vector of costate variables ***λ***(*t*) = (*λ*_1_(*t*), *λ*_2_(*t*), …, *λ*_*n*_(*t*)) and define a Hamiltonian

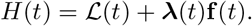

The costate variables can be obtained from the necessary conditions

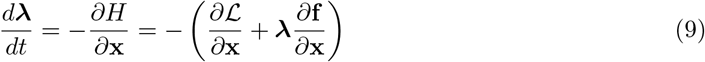

and the transversality condition

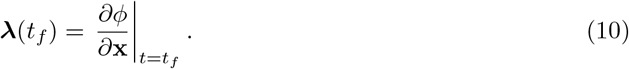

Pontryagin’s maximum (minimum) principle [23] states that the cost function is minimised when the control, together with the corresponding state and costate, minimise *H*(*t*) for all *t* ∈ [*t*_0_, *t*_*f*_]. In general this is not directly solvable, as **x** and ***λ*** must be obtained numerically for a given *u*. We use the following approach, where at each step *t* ∈ [*t*_0_, *t*_*f*_]:

**Algorithm 1**

1. Select an initial value for *u*(*t*).
2. Solve the boundary value problem given by the state Equations (1) and (2) for **x**(*t*).
3. Solve the boundary value problem given by the costate Equations (9) and (10) for ***λ***(*t*).
4. Find *u*^∗^(*t*) which minimises *H*(*t*) for the given **x**(*t*) and ***λ***(*t*).
5. Update *u*(*t*) based on a combination of the current value and *u*^∗^(*t*).
6. Check the specified convergence condition; if not met, go to step 2.

We use the initial control value *u*(*t*) = 0 in all cases. The state equations **f** (**x**, *u*) are given by Equations (4-6), with **x** = (*A, P, N*). In all cases the initial state is a stable equilibrium with cancer present and no control (see below). We solve the boundary value problems using the fourth order Runge-Kutta method and time step 0.001. In step 3, the boundary value is specified at time *t*_*f*_, so the solution is obtained working backwards in time. The details of the costate equations and of steps 4, 5 and 6 depend on the cost function and the corresponding optimal control form, as discussed in Section 1.1. We consider the two cases separately below. We use the convergence condition |*u*_*new*_ − *u*_*old*_|*/*|*u*_*new*_| *<* 10^−3^.

A limitation of the optimal control methodology is that results are valid only for the specified time window, while we are typically interested in the cost of treatment over an indefinite period. This includes cases where ongoing treatment is required. If the model parameters do not permit permanent control of the cancer, the optimal control may include an artefact late in the time window caused by the artificial end-point. For this reason all optimal controls were obtained using a time window [0, 200], and only the period [0, 100] or [0, 50] was shown. All costs and other statistics shown or plotted were recalculated on this interval.

### 4.2 Continuous control

We first consider the quadratic cost function ℒ = *u*^2^ + (*P* + *N*)^2^ (Equation 7), which will typically correspond to continuous optimal control solutions. Using Equation 9 with 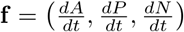 given by (4-6), we obtain the costate equations

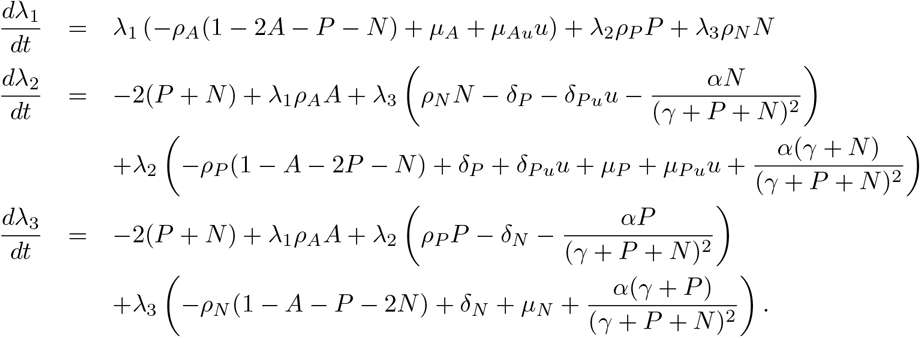

The end-state cost *ϕ* in the cost function (3) is set to zero, so the boundary condition (10) gives ***λ***(*t*_*f*_) = 0. We next find *u*^∗^(*t*) that minimises *H* for all *t* ∈ [*t*_0_, *t*_*f*_]. Note that

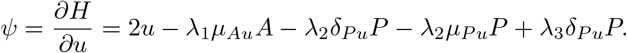

Setting *ψ*(*t*) = 0 gives

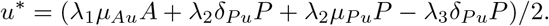

Using *u*^∗^ as the updated value of *u* will not in general allow for convergence, and so the updated value of *u* is then taken to be a linear combination of the current value and *u*^∗^,

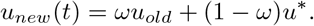

The parameter *ω* ∈ [0, 1) can be increased as required to achieve convergence; we used *ω* = 0.9 in all cases, with convergence in less than 100 steps.

### 4.3 Bang-bang control

We now consider the linear cost function ℒ = *u* + *P* + *N* (Equation 8). The costate equations differ only slightly from the continuous control case, with the term −2(*P* + *N*) replaced by −1 in the equations for 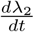 and 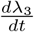. The costate boundary condition is again ***λ***(*t*_*f*_) = 0. However, a substantially different approach is required for the optimality condition. Note that

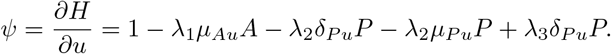

Since *ψ*(*t*) is not a function of *u*, the natural approach to minimising *H* is to increase *u* where *ψ*(*t*) *<* 0, and decrease *u* where *ψ*(*t*) *>* 0, while limiting the change in *u* in each step to allow convergence given the indirect effect of *u* on *H* via **x** and ***λ***. However, this will generally result in *u* diverging towards ±∞ at some values of *t*, and so it is necessary to impose bounds *u*_0_ *< u*(*t*) *< u*_1_. Such bounds are typically consistent with the real world system being modelled. In our model, the drug level cannot be negative and there will be an upper bound on the safe and effective drug dose; we use the range [*u*_0_, *u*_1_] = [0, 1]. In the approach adopted from [27], we set *u*^∗^(*t*) = *u*_0_ for *ψ*(*t*) *>* 0, and *u*^∗^(*t*) = *u*_1_ for *ψ*(*t*) *<* 0. We then set *u*_*new*_(*t*) = *ωu*_*old*_ + (1 − *ω*)*u*^∗^, as for the continuous control. In some cases we found that this algorithm converged to a solution in which *u*(*t*) is equal to either 0 or 1 for each *t* ∈ [*t*_0_, *t*_*f*_], the expected form of a bang-bang control. But note that while *u*^∗^(*t*) has this form at each time step by definition, the update method using a weighted average means that *u*(*t*) does not have this form at each step of the iteration. In fact, we found that for some parameter cases we could not meet the formal convergence criterion for any choice of *ω*, yet the control appeared to approximately converge to a form in which *u*(*t*) attains a value intermediate between 0 and 1 for a range of *t*. The algorithm given above cannot achieve convergence to such a control, since *u*^∗^(*t*) will be equal to 0 or 1 for any given iteration and value of *t*. In order to achieve convergence to optimal control solutions of this type, we adopted the following convergence strategy:

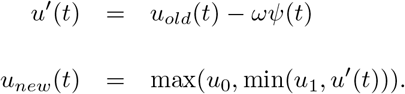

The convergence parameter *ω >* 0 is not comparable to the parameter in the previous method. Lower values give slower but more reliable convergence. We used *ω* ∈ {0.02, 0.01, 0.005}. Convergence speed was found to be substantially lower than for continuous controls even for simple bang-bang controls using the original algorithm. For the modified algorithm we added an additional secondary convergence condition, calculated every 1000 iterations: |*u*_*t*_ − *u*_*t*−1000_|*/*|*u*_*t*_| *<* 10^−2^. For obtaining controls including periods of intermediate control values using the modified method, convergence speed was substantially slower again, with up to 10000 iterations required for convergence, and up to 60000 iterations were required for more complex control solutions.

### 4.4 Equilibria

All optimal control calculations use an initial state **x**(*t*_0_) in which cancer is present and the system is in stable equilibrium prior to application of the control. We do not have an exact expression for this steady state, so we first find the steady state solution with *N, P >* 0 for the restricted model with *α* = 0 (no immune response). We then run the full model simulation, using the fourth order Runge-Kutta method and time step 0.001 as in the optimal control algorithm, until the absolute single time step change in each state variable is less than the minimum floating point difference (2^−53^).

In the following we provide the required derivation for the steady states with *α* = 0. We also show that the physically realisable steady states for this model conform to one of two cases: (1) there is exactly one steady state, which has *N* = *P* = 0 and is stable; (2) there are two steady states, one with *N* = *P* = 0 which is unstable, and one with *P, N >* 0 which is stable. In particular, this shows that the Michaelis-Menten immune response term is necessary in order to allow for cases in which cancer is present and persistent, but can be permanently controlled by a finite period of drug treatment. Note that in the neighbourhood of *P* = *N* = 0, the full model is equivalent to a model with *α* = 0 and modified cancer exit rates 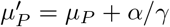 and 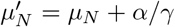. Using the method below we can thus determine exactly when the *N* = *P* = 0 equilibrium is stable. However, in the full model a stable *P* = *N* = 0 equilibrium does not exclude a stable equilibrium with *N, P >* 0, due to the reduction in the immune component of the exit rates as *P* + *N* increases.

#### 4.4.1 Equilibria with zero immune response

To obtain the steady states of our model with no control and in the absence of the immune response, we let *C* = 1 − *A* − *P* − *N* and set 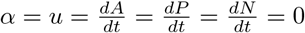 in (4-6) to give

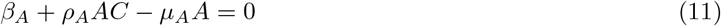

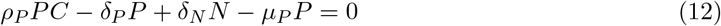

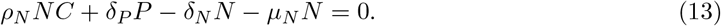

For physical (non-negative) *N* and *P*, we have *A* ≤ 1 − *C*. With (11) this gives the condition 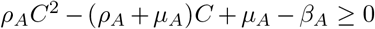. The larger zero is greater than 1 and hence non-physical, thus

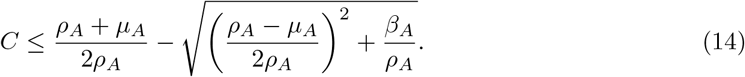

For any *C* satisfying this constraint, we obtain *A* by rearranging Equation 11 to give *A* = *β*_*A*_*/*(*µ*_*A*_ − *ρ*_*A*_*C*). We always have a non-cancer steady state in which *P* = *N* = 0, where *C* is equal to the bound in (14) and *A* = 1 − *C*. By (12) and (13), *N* = 0 ⇔ *P* = 0, so it remains only to consider the case *N >* 0 and *P >* 0. From (12) and (13) we have

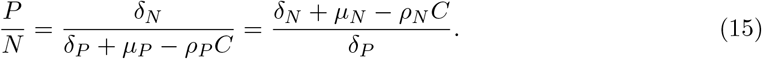

This gives a quadratic in *C*. The larger solution is greater than 1, and hence the only potentially physical solution is

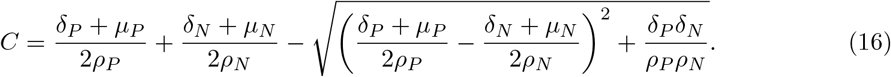

We see that there are two cases. There is always a non-cancerous steady state. If the expression for *C* in (16) satisfies condition (14) without exact equality, and gives a positive value for the ratio *P/N* in (15), then *A* can then be obtained from (11), and the cancerous cells *N* + *P* = 1 − *A* – *C* can be apportioned according to (15) to give exactly one additional physical solution.

We next consider the stability of these equilibria. The Jacobian for the system when *α* = *u* = 0 is

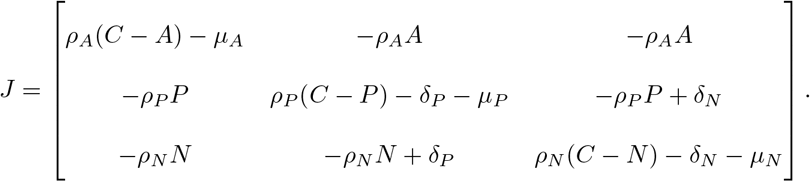

For the non-cancerous fixed point (*A, P, N*) = (*A*_0_, 0, 0), the characteristic equation det(*J* − *λI*) = 0 is

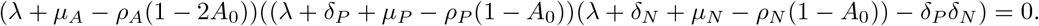

The equilibrium is stable if and only if all solutions have negative real part. The first term gives the solution *λ* = *ρ*_*A*_(1 − 2*A*_0_) − *µ*_*A*_ *< β*_*A*_ + *ρ*_*A*_*A*_0_(1 − *A*_0_) − *µ*_*A*_*A*_0_, which is negative by (11). This leaves the solutions to the quadratic

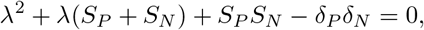

where *S*_*P*_ = *δ*_*P*_ + *µ*_*P*_ − *ρ*_*P*_ (1 − *A*_0_) and *S*_*N*_ = *δ*_*N*_ + *µ*_*N*_ − *ρ*_*N*_ (1 − *A*_0_). Thus by the Routh–Hurwitz stability criterion, all solutions will have a negative real part if and only if *S*_*P*_ + *S*_*N*_ *>* 0 and *S*_*P*_ *S*_*N*_ *> δ*_*P*_ *δ*_*N*_.

Let *C* = *C*_1_ be the solution of (16), and define 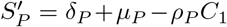 and 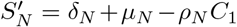. We saw above that there exists a physical equilibrium with non-zero cancer exactly when *C*_1_ *<* 1 − *A*_0_ and the *P/N* ratio given by (15) for *C* = *C*_1_ is positive (note that the equality in (15) holds by the definition of *C*_1_). This implies that 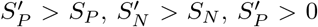 and 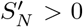, and hence either 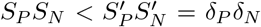 or else both *S*_*P*_ and *S*_*N*_ are negative. Thus (*A*_0_, 0, 0) is unstable.

Conversely, if (*A*_0_, 0, 0) is stable then 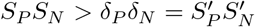 and *S*_*P*_, *S*_*N*_ *>* 0. This implies that either *C*_1_ *>* 1 − *A*_0_ or both 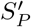 and 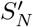 are negative. Thus there is no real physical solution with *N,P>*0.

We now suppose that there exists a steady state (*A*_1_, *P*_1_, *N*_1_) with *A*_1_, *P*_1_, *N*_1_ *>* 0, and consider stability at this point. Let *C*_1_ = 1 − *A*_1_ − *P*_1_ − *N*_1_ and *R* = *P*_1_*/N*_1_ *>* 0. By (11) and (15) we have *µ*_*A*_ − *ρ*_*A*_*C*_1_ = *β*_*A*_*/A*_1_, *δ*_*P*_ + *µ*_*P*_ − *ρ*_*P*_ *C*_1_ = *δ*_*N*_ */R*, and *δ*_*N*_ + *µ*_*N*_ − *ρ*_*N*_ *C*_1_ = *Rδ*_*P*_, giving

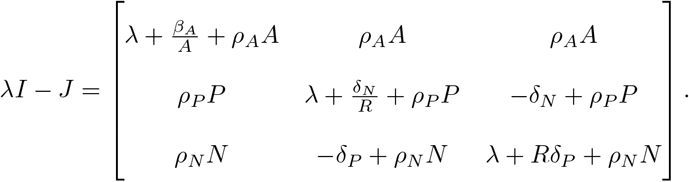

Noting the cancellation of all terms containing (*ρ*_*A*_*A*)(*ρ*_*P*_ *P*), (*ρ*_*A*_*A*)(*ρ*_*N*_ *N*), or (*ρ*_*P*_ *P*)(*ρ*_*N*_ *N*) in det(*λI* − *J*), we obtain the characteristic equation

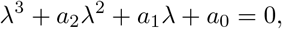

where

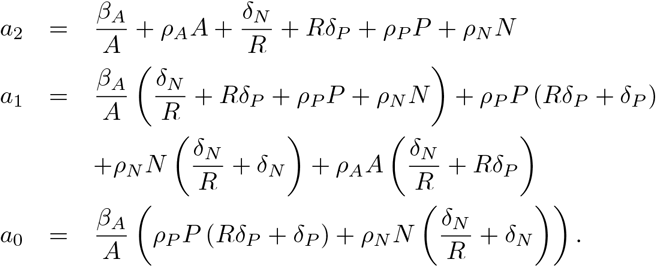

Since all parameters and state variables are positive, we observe that *a*_2_, *a*_1_, *a*_0_ *>* 0 and *a*_2_*a*_1_ *> a*_0_. Thus by the Routh-Hurwitz criterion all solutions have a negative real part, and the fixed point (*A*_1_, *P*_1_, *N*_1_) is stable.

Thus we have shown that there is either a single, stable fixed point with *P* = *N* = 0, or else this fixed point is unstable and there is a second, stable fixed point with *P, N >* 0.

## 5 Data availability statement

There are no primary data in the paper; all code is available on a GitHub repository at https://github.com/jameslefevreoptimal-control/tree/main.

## References

[1] Marc S Raab, Klaus Podar, Iris Breitkreutz, Paul G Richardson, and Kenneth C Anderson. Multiple myeloma. The Lancet, 374(9686):324–339, jul 2009. doi: 10.1016/s0140-6736(09)60221-x.

[2] Kazuhito Suzuki, Kaichi Nishiwaki, and Shingo Yano. Treatment strategy for multiple myeloma to improve immunological environment and maintain MRD negativity. Cancers, 13(19):4867, sep 2021. doi: 10.3390/cancers13194867.

[3] Torben Plesner and Jakub Krejcik. Daratumumab for the treatment of multiple myeloma. Frontiers in Immunology, 9, jun 2018. doi: 10.3389/fimmu.2018.01228.

[4] Jakub Krejcik, Tineke Casneuf, Inger S. Nijhof, Bie Verbist, Jaime Bald, Torben Plesner, Khaja Syed, Kevin Liu, Niels W. C. J. van de Donk, Brendan M. Weiss, Tahamtan Ahmadi, Henk M. Lokhorst, Tuna Mutis, and A. Kate Sasser. Daratumumab depletes CD38+ immune regulatory cells, promotes t-cell expansion, and skews t-cell repertoire in multiple myeloma. Blood, 128(3):384–394, jul 2016. doi: 10.1182/blood-2015-12-687749.

[5] Yu-Tzu Tai, Myles Dillon, Weihua Song, Merav Leiba, Xian-Feng Li, Peter Burger, Alfred I. Lee, Klaus Podar, Teru Hideshima, Audie G. Rice, Anne van Abbema, Lynne Jesaitis, Ingrid Caras, Debbie Law, Edie Weller, Wanling Xie, Paul Richardson, Nikhil C. Munshi, Claire Mathiot, Hervé Avet-Loiseau, Daniel E. H. Afar, and Kenneth C. Anderson. Anti-cs1 humanized monoclonal antibody huluc63 inhibits myeloma cell adhesion and induces antibody-dependent cellular cytotoxicity in the bone marrow milieu. Blood, 112(4):1329–1337, August 2008. ISSN 1528-0020. doi: 10.1182/blood-2007-08-107292.

[6] Anthony Markham. Elotuzumab: First global approval. Drugs, 76(3):397–403, January 2016. ISSN 1179-1950. doi: 10.1007/s40265-016-0540-0.

[7] X. Armoiry, G. Aulagner, and T. Facon. Lenalidomide in the treatment of multiple myeloma: a review. Journal of Clinical Pharmacy and Therapeutics, 33(3):219–226, June 2008. ISSN 1365-2710. doi: 10.1111/j.1365-2710.2008.00920.x.

[8] Sagar Lonial, Meletios Dimopoulos, Antonio Palumbo, Darrell White, Sebastian Grosicki, Ivan Spicka, Adam Walter-Croneck, Philippe Moreau, Maria-Victoria Mateos, Hila Magen, Andrew Belch, Donna Reece, Meral Beksac, Andrew Spencer, Heather Oakervee, Robert Z. Orlowski, Masafumi Taniwaki, Christoph Röllig, Hermann Einsele, Ka Lung Wu, Anil Singhal, Jesus San-Miguel, Morio Matsumoto, Jessica Katz, Eric Bleickardt, Valerie Poulart, Kenneth C. Anderson, and Paul Richardson. Elotuzumab therapy for relapsed or refractory multiple myeloma. New England Journal of Medicine, 373(7):621–631, August 2015. ISSN 1533-4406. doi: 10.1056/nejmoa1505654.

[9] Ilaria Saltarella, Vanessa Desantis, Assunta Melaccio, Antonio Giovanni Solimando, Aurelia Lamanuzzi, Roberto Ria, Clelia Tiziana Storlazzi, Maria Addolorata Mariggiò, Angelo Vacca, and Maria Antonia Frassanito. Mechanisms of resistance to anti-CD38 daratumumab in multiple myeloma. Cells, 9(1):167, jan 2020. doi: 10.3390/cells9010167.

[10] Harold C. Sullivan, Christian Gerner-Smidt, Ajay K. Nooka, Connie M. Arthur, Louisa Thompson, Amanda Mener, Seema R. Patel, Marianne Yee, Ross M. Fasano, Cassandra D. Josephson, Richard M. Kaufman, John D. Roback, Sagar Lonial, and Sean R. Stowell. Daratumumab (anti-CD38) induces loss of CD38 on red blood cells. Blood, 129(22):3033–3037, jun 2017. doi: 10.1182/blood-2016-11-749432.

[11] Philippe Moreau, Cyrille Hulin, Aurore Perrot, Bertrand Arnulf, Karim Belhadj, Lotfi Benboubker, Marie C Béné, Sonja Zweegman, Hélène Caillon, Denis Caillot, Jill Corre, Michel Delforge, Thomas Dejoie, Chantal Doyen, Thierry Facon, Cécile Sonntag, Jean Fontan, Mohamad Mohty, Kon-Siong Jie, Lionel Karlin, Frédérique Kuhnowski, Jérôme Lambert, Xavier Leleu, Margaret Macro, Frédérique Orsini-Piocelle, Murielle Roussel, Anne-Marie Stoppa, Niels W C J van de Donk, Soraya Wuillème, Annemiek Broijl, Cyrille Touzeau, Mourad Tiab, Jean-Pierre Marolleau, Nathalie Meuleman, Marie-Christiane Vekemans, Matthijs Westerman, Saskia K Klein, Mark-David Levin, Fritz Offner, Martine Escoffre-Barbe, Jean-Richard Eveillard, Réda Garidi, Tahamtan Ahmadi, Maria Krevvata, Ke Zhang, Carla de Boer, Sanjay Vara, Tobias Kampfenkel, Veronique Vanquickelberghe, Jessica Vermeulen, Hervé Avet-Loiseau, and Pieter Sonneveld. Maintenance with daratumumab or observation following treatment with bortezomib, thalidomide, and dexamethasone with or without daratumumab and autologous stem-cell transplant in patients with newly diagnosed multiple myeloma (CAS-SIOPEIA): an open-label, randomised, phase 3 trial. The Lancet Oncology, 22(10):1378–1390, oct 2021. doi: 10.1016/s1470-2045(21)00428-9.

[12] Philip Holmes. Poincaré, celectial mechanics, dynamical-systems theory and “chaos”. Physics Reports (review section of Physics Letters) 193, No. 3, 1990.

[13] Pierre-François Verhulst. Notice sur la loi que la population poursuit dans son accroissement. Correspondance Mathématique et Physique, 10:113–121, 1838.

[14] Pierre-François Verhulst. Recherches mathématiques sur la loi d’accroissement de la population [mathematical researches into the law of population growth increase]. Nouveaux Mémoires de l’Académie Royale des Sciences et Belles-Lettres de Bruxelles, 18(8), 1845.

[15] Alfred J. Lotka. Contribution to the theory of periodic reactions. The Journal of Physical Chemistry, 14(3):271–274, mar 1910. doi: 10.1021/j150111a004.

[16] Narendra S. Goal, Samaresh C. Maitra, and Elliott W. Montroll. On the volterra and other nonlinear models of interacting populations. Reviews of Modern Physics, 43(2):231–276, apr 1971. doi: 10.1103/revmodphys.43.231.

[17] Abicumaran Uthamacumaran. A review of dynamical systems approaches for the detection of chaotic attractors in cancer networks. Patterns, 2(4):100226, apr 2021. doi: 10.1016/j.patter.2021.100226.

[18] Odelaisy León-Triana, Soukaina Sabir, Gabriel F. Calvo, Juan Belmonte-Beitia, Salvador Chulián, Álvaro Martínez-Rubio, María Rosa, Antonio Pérez-Martínez, Manuel Ramirez-Orellana, and Víctor M. Pérez-García. CAR t cell therapy in b-cell acute lymphoblastic leukaemia: Insights from mathematical models. Communications in Nonlinear Science and Numerical Simulation, 94:105570, mar 2021. doi: 10.1016/j.cnsns.2020.105570.

[19] Manju Agarwal and Archana S. Bhadauria. A generalised prey-predator type model of immunogenic cancer with the effect of immunotherapy. International Journal of Engineering, Science and Technology, 5(1):66–84, mar 2018. doi: 10.4314/ijest.v5i1.6.

[20] Robert C. Sterner and Rosalie M. Sterner. CAR-t cell therapy: current limitations and potential strategies. Blood Cancer Journal, 11(4), apr 2021. doi: 10.1038/s41408-021-00459-7.

[21] Irina Kareva, Kimberly A. Luddy, Cliona O’Farrelly, Robert A. Gatenby, and Joel S. Brown. Predator-prey in tumor-immune interactions: A wrong model or just an incomplete one? Frontiers in Immunology, 12, aug 2021. doi: 10.3389/fimmu.2021.668221.

[22] Phineas T. Hamilton, Bradley R. Anholt, and Brad H. Nelson. Tumour immunotherapy: lessons from predator–prey theory. Nature Reviews Immunology, may 2022. doi: 10.1038/s41577-022-00719-y.

[23] L.S. Pontryagin, V.G. Boltyanskii, R.V. Gamkrelidze, and E.F. Mischenko. The mathematical theory of optimal processes [english translation]. 1962.

[24] N.L. Grigorenko, E.N. Khailov, E.V. Grigorieva, and A.D. Klimenkova. Optimal strategies for car-t therapy for the treatment of leukemia in the lotka-volterra predator-prey model. 27 (3):43–58, sep 2021. doi: 10.21538/0134-4889-2021-27-3-43-58.

[25] Pariya Khalili and Ramin Vatankhah. Optimal control design for drug delivery of immunotherapy in chemoimmunotherapy treatment. Computer Methods and Programs in Biomedicine, 229:107248, feb 2023. doi: 10.1016/j.cmpb.2022.107248.

[26] Helena L. Crowell, Adam L. MacLean, and Michael P.H. Stumpf. Feedback mechanisms control coexistence in a stem cell model of acute myeloid leukaemia. Journal of Theoretical Biology, 401:43–53, jul 2016. doi: 10.1016/j.jtbi.2016.04.002.

[27] Jesse A. Sharp, Alexander P Browning, Tarunendu Mapder, Kevin Burrage, and Matthew J Simpson. Optimal control of acute myeloid leukaemia. Journal of Theoretical Biology, 470: 30–42, jun 2019. doi: 10.1016/j.jtbi.2019.03.006.

